# Attention and microsaccades: do attention shifts trigger new microsaccades or only bias ongoing microsaccades?

**DOI:** 10.1101/2023.09.28.559945

**Authors:** Baiwei Liu, Zampeta-Sofia Alexopoulou, Freek van Ede

## Abstract

Brain circuitry that controls where we look also contributes to attentional focusing of visual contents outside of current fixation or contents held within the spatial layout of working memory. A behavioural manifestation of the contribution of oculomotor brain circuitry to selective attention comes from modulations in microsaccade direction that accompany attention shifts. Here, we address whether such modulations come about because attention itself triggers new microsaccades or whether, instead, shifts in attention only bias the direction of ongoing microsaccades – i.e., naturally occurring microsaccades that would have been made whether or not attention was also shifted. We utilised an internal-selective-attention task that has recently been shown to yield clear spatial microsaccade modulations and compared microsaccade rates following colour retrocues that were matched for sensory input, but differed in whether they invited an attention shift or not. If shifts in attention trigger new microsaccades then we would expect more microsaccades following attention-directing cues than following neutral cues. In contrast, we found no evidence for an increase in overall microsaccade rate following attention-directing cues, despite observing robust modulations in microsaccade direction. This implies that shifting attention biases the direction of ongoing microsaccades without changing the probability that a microsaccade will occur. These findings provide relevant context for complementary and future work delineating the links between attention, microsaccades, and upstream oculomotor brain circuitry, such as by helping to explain why microsaccades and attention shifts are often correlated but not obligatorily linked.

## INTRODUCTION

Goal-driven visual attention can be directed not only overtly – by looking directly at task-relevant visual information – but also covertly, by attending to relevant information outside of current fixation (e.g., (Carrasco, 2011; Desimone & Duncan, 1995)) or to information held internally within working memory (e.g., (Griffin & Nobre, 2003; Souza & Oberauer, 2016; van Ede & Nobre, 2023)). Ample prior research has demonstrated that brain circuitry that is involved in overt eye-movement control also contributes to the deployment of covert visual-spatial attention (e.g., (Krauzlis et al., 2013; Kustov & Robinson, 1996; Moore et al., 2003; Nobre et al., 2000; Rizzolatti et al., 1987)).

One particular example of this contribution comes from the study of microsaccades, a class of fixational eye-movements that occur even during attempted fixation (e.g., (Engbert, 2006; Hafed et al., 2015; Martinez-Conde et al., 2004; Rolfs, 2009; Rucci & Poletti, 2015)). In particular, it has been demonstrated how the direction of fixational microsaccades can be modulated by the deployment of spatial attention to peripherally attended locations, in the absence of large eye-movements to these locations ((Corneil & Munoz, 2014; Engbert, 2012; Engbert & Kliegl, 2003; Fernández et al., 2023; Hafed et al., 2011; Hafed & Clark, 2002; Laubrock et al., 2005; Lowet et al., 2018; Pastukhov & Braun, 2010; Xue et al., 2020); but see also (Horowitz et al., 2007; Tse et al., 2002; Willett & Mayo, 2023) to which we return in our discussion). Building on this earlier work, we recently uncovered similar microsaccade biases when directing attention to memorised visual contents held within the spatial lay-out of working memory (e.g., (de Vries & van Ede, 2023; Liu et al., 2022; van Ede et al., 2019, 2020, 2021)). This provides a useful model system for studying the link between microsaccades and attention because in this set-up there is no incentive for large eye-movements as the objects of attention are in mind.

When considering the link between microsaccades and attention, a relevant question is whether shifting attention will itself trigger new microsaccades or whether, instead, shifting attention merely biases the direction of *ongoing* microsaccades – i.e., naturally occurring microsaccades that would have been made whether or not attention was also shifted.

These alternative scenarios have often remained tacit in the literature. Yet, disambiguating these scenarios is likely to be informative for our understanding of the links between attention, microsaccades, and upstream oculomotor brain circuitry. Foremost, the answer to our question may help delineate the probabilistic (e.g., (Horowitz et al., 2007; Liu et al., 2022; Yu et al., 2022)) versus deterministic (e.g., (Lowet et al., 2018)) nature of the link between microsaccades and covert attention shifts. If attention only biases the direction of ongoing microsaccades, without triggering new microsaccades, then this would explain a probabilistic link whereby attention can also be shifted without a concomitant microsaccade (corroborating the findings reported in (Liu et al., 2022; Yu et al., 2022)). In addition, our findings may guide future neurophysiological studies targeting the role of upstream oculomotor brain circuitry, such as the superior colliculus – a brain structure implicated in both selective covert attention (Krauzlis et al., 2013; Lovejoy & Krauzlis, 2010; Muller et al., 2005) and microsaccade generation (Hafed et al., 2009; Hafed & Krauzlis, 2012). For example, based on our results, we may derive distinct hypotheses that attention adds new activity to the pool of superior-colliculus neurons (triggering new microsaccades) or instead mainly acts by biasing the balance of neuronal activity (yielding only a bias in the *direction* but not the *rate* of microsaccades).

To address whether attention shifts add new microsaccades or bias ongoing ones, we compared microsaccade rates following attentional cues that were carefully matched for sensory input, but that differed whether they invited an attention shift or not. We note how the comparison between informative (attention-directing) and neutral cues was also available in (Engbert & Kliegl, 2003; Laubrock et al., 2005) though in these studies the spatial biasing by voluntary attention itself was weak and the cues were not perfectly matched, making it hard to address the question that we put central here. Our logic was straightforward: if voluntary attention shifts trigger new microsaccades – that account for the observed spatial modulation in microsaccade direction – then overall microsaccade rates should be higher following attention-directing than following neutral cues. In contrast, if attention merely biases ongoing microsaccades, then we should see a biasing effect on microsaccade direction *without* a concomitant increase in overall microsaccade rate.

## METHODS

### Ethics

Experimental procedures were reviewed and approved by the local Ethics Committee at the Vrije Universiteit Amsterdam. Each participant provided written informed consent before participation and was reimbursed 10 euros/hour.

### Participants

Twenty-five healthy human volunteers participated in the study (age range: 18 - 44; 5 male and 20 female; 25 right-handed; 5 corrected-to-normal vision: 1 glasses and 4 lenses). Sample size of 25 was determined a-priori based on previous publications from the lab with similar experimental designs that relied on the same outcome measure (e.g., (van Ede et al., 2019, 2020, 2021)). One participant was excluded for all analyses due to chance-level performance.

### Stimuli and procedure

To investigate microsaccade modulations by voluntary shifts of attention, we employed an internal selective-attention task (**Fig. 1**) for which we previously established robust spatial modulations in microsaccades (see e.g., (Liu et al., 2022; van Ede et al., 2019)). In short, participants encoded two visual items into working memory in order to compare the orientation of either memory item to an upcoming test stimulus. In a random half of the trials, a retrocue presented during the retention interval informed which memory item would become tested by briefly changing the colour of the central fixation marker to match the colour of the target memory item (attention-directing cue). In the other half of the trials, we also presented a colour cue, but this time cue colour did not match either item in memory, and hence did not invite a shift of attention to either memory content (neutral cue).

**Figure 1.**
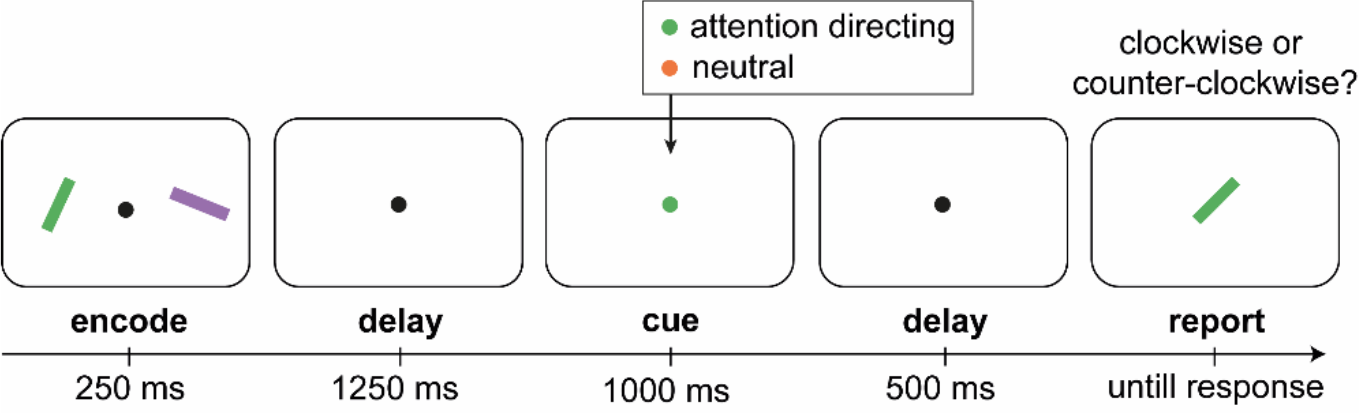
Internal selective attention task with attention-directing and neutral colour retrocues. Participants encoded two visual items into working memory in order to later compare the orientation of one of the items to a test stimulus that was tilted 10 or 20 degrees clockwise or counter-clockwise relative to the colour-matching memory item. During the delay, the colour of the central fixation dot changed colour serving as a cue. In a random half of the trials, the retrocue matched the colour of either item in working memory, informing with 100% reliability that this item would become tested. In the other trials, the cue also involved a colour change, but this time the colour did not match either item in working memory. Colours of memory items and cues were counterbalanced, such that a physically identical colour cue would be attention directing in some trials while neutral in other trials.

Each trial began with a brief (250 ms) encoding display in which two bars (size: 2° × 0.4° visual angle) appeared at 5° to the left and right of the fixation. After an initial retention delay of 1250 ms, the fixation dot (0.07° radius) changed colour for 1000 ms serving as a retrocue that prompted participants to select the colour-matching target item in memory. After another retention delay of 500 ms, the test display appeared in which a target bar appeared at the center of the screen. The target bar matched the colour of the target memory item but was rotated between 10 to 20 degrees clockwise or counter-clockwise from its original orientation. Participants were required to report the tilt offset of the test stimulus using the keyboard (‘j’ for clockwise, ‘f’ for counter-clockwise). Participants received feedback immediately after the response by a number (“0” for wrong, or “1” for correct) appearing for 250 ms slightly above the fixation dot. After the feedback, inter-trial intervals were randomly drawn between 500 and 1000 ms.

In the experiment, bars could be four potential colours: green (RGB: 133, 194, 18), purple (RGB: 197, 21, 234), orange (RGB: 234, 74, 21), and blue (RGB: 21, 165, 234]). For each participant, bars were always chosen from a random subset of three of these colours. In each encoding display, bars were randomly assigned two distinct colours from the available colour pool and two distinct orientations ranging from 0° to 180° with a minimum difference of 20° between each other.

To address our central question whether attention triggers additional microsaccades, we included both attention-directing and neutral retrocues. In half the trials, the retrocue colour matched either memory item, inviting a shift of attention to the to-be-tested memory content. Participants were encouraged to use these informative, attention-directing retrocues that were 100% valid. In the other half of the trials, the retrocue was drawn from either colour that was *not* in the encoding display, thus not inviting a shift of attention among the contents of working memory. In these trials, participants would know which memory item was the target memory item only upon the presentation of the coloured test stimulus.

In total, the study consisted of 2 sessions, each containing 5 blocks of 48 trials, resulting in a total of 480 trials. Both conditions (attention-directing cues and neutral cues) were randomly intermixed within each block as were attention directing cues to left and right memory items, resulting in 240 attention-directing trials (120 directing attention to the left memory item, 120 directing attention to the right memory item), and 240 neutral trials. At the start of the experiment, participants practiced the task for 48 trials. We did not include practice trials in our analyses.

### Eye-tracking acquisition and pre-processing

Using an EyeLink 1000 with a sampling rate of 1000 Hz, we continuously tracked gaze along the horizontal and vertical axes from the right eye. The eye tracker was placed ∼5 cm in front of the monitor and ∼65 cm away from the eyes. Before recording, we calibrated the eye tracker through the built-in calibration and validation protocols from the EyeLink software. Gaze data was originally recorded in .edf format and was converted to .asc format to be further analyzed after recording.

We analysed the data in Matlab with help of the Fieldtrip analysis toolbox (Oostenveld et al., 2011) and custom code. To clean the data from blinks, we marked blinks by detecting clusters of zeros in the time-series eye data. To eliminate residual blink artifacts, all data from 100 ms before to 100 ms after the detected blink clusters were set to Not-a-Number (NaN) and thereby ignored in further analysis. After blink removal, data were epoched relative to retrocue onset.

### Saccade detection

To detect saccades, we employed a velocity-based method that we established previously (Liu et al., 2022), and that builds on other established velocity-based methods for microsaccade detection (e.g., (Engbert & Kliegl, 2003)). Since the items in the current experiment were always horizontally arranged (i.e., left and right), our current analyses focused exclusively on the horizontal channel of the eye data. Note that although we only use horizontal data to detect saccades, we previously confirmed the validity and sensitivity of this approach for our task-setup by comparing this method to a well-established method (as described in (Engbert & Kliegl, 2003)) that considered both horizontal and vertical gaze (see (Liu et al., 2022) for the relevant comparison).

We first calculated the gaze velocity by taking the distance between temporally successive gaze positions. Then, to reduce noise, we smoothed velocity in the temporal dimension with a Gaussian-weighted moving average filter with a 7-ms sliding window (using the built-in function “smoothdata” in MATLAB). We then identified the first sample when the velocity exceeded a trial-based threshold of 5 times the median velocity as the onset of a saccade. To avoid counting the same saccade multiple times, we imposed a minimum delay of 100 ms between successive saccades. Saccade magnitude and direction were calculated by estimating the difference between pre-saccade gaze position (−50 to 0 ms before threshold crossing) vs. the post-saccade gaze position (50 to 100 ms after threshold crossing). Finally, depending on saccade direction (left/right) and the side of the cued memory item (left/right), we labeled every detected saccades as “toward” or “away”.

After identifying and labelling the saccades, we quantified the time courses of saccade rates (in Hz) using a sliding time window of 50 ms, advanced in steps of 1 ms. To map the size of the modulated saccades (without setting an arbitrary saccade-size threshold), we additionally decomposed saccade rates into a time-size representation (as in (Liu et al., 2022)), showing the time courses of saccade rates, as a function of the saccade size. For saccade-size sorting, we used successive magnitude bins of 0.5 visual degrees in steps of 0.05 visual degree.

To directly quantify the number of saccades during the attentional window of interest we averaged saccade rates in the 200-600 ms window after cue onset. This window was set a-priori based on our prior study that revealed this to be the critical window after cue onset in which we found more microsaccades toward vs. away from the memorized location of the cued memory target (see (Liu et al., 2022)).

### Statistical analysis

To evaluate the reliability of statistical patterns we observed in the time-series data, we employed a cluster-based permutation approach (Maris & Oostenveld, 2007). This method is ideal for evaluating significance while circumventing the problem of multiple comparisons.

We first acquired a permutation distribution of the largest cluster size by randomly permuting the trial-average data at the participant level 10,000 times and identifying the size of the largest clusters after each permutation. To obtain the probability (P value) of the clusters observed in the original data, we calculated the proportion of permutations for which the size of the largest cluster after permutation was larger than the size of the observed cluster in the original, non-permuted data. The permutation analysis was conducted using Fieldtrip with default clustering settings. That is, after a mass univariate t-test at a two-sided alpha level of 0.05, we identified and grouped the adjacent same-signed data points that were significant and then defined cluster size as the sum of all t-values in the cluster.

In addition to the cluster-based permutation approach that considered the full time range, we also extracted the data over the pre-defined 200-600 ms window after cue onset, and compared the relevant conditions using paired-sample t-tests.

### Data and code availability

Data and analysis code will be made publicly available before publication.

## RESULTS

Human volunteers performed a selective-attention task in which attention was directed to one of two visual representations in working memory (**Fig. 1**). In half of the trials, a central colour cue directed attention to the colour-matching visual item in working memory (attention-directing cues). In the other half of the trials, we also presented a colour cue but this time the cue did not match either memory item and therefore did not invite a shift of attention (neutral cues). This served as the critical control condition to assess whether shifting attention triggered new microsaccades: in both cases a central colour cue appeared but only in the former condition a goal-directed shift of attention could be made.

As a roadmap to our results, we first report behavioural performance to confirm that participants used the cue when it invited a shift of attention to the target memory item. We then outline the spatial modulation of microsaccade direction when cues directed attention to either the left or right memory item. Having established the above, we finally turn to our key question whether this spatial microsaccade modulation is driven by the addition of new, attention-driven, microsaccades or, instead, by a biasing of ongoing microsaccades that would have been made anyway. For this, we compared overall microsaccade rate between trials with attention-directing cues versus neutral cues. The logic is straightforward: if attention adds new microsaccades then overall rate should increase following attention-directing compared to neutral cues. In contrast, if attention merely biases the direction of ongoing microsaccades then we should observe similar rates following attention-directing and neutral cues.

### Informative attentional cues during working memory improve ensuing memory-guided behaviour

Before turning to our main eye-movement results, we first turn to the behavioural performance in our task, to confirm that informative (attention-directing) cues were used by participants (c.f. (Griffin & Nobre, 2003)). As shown in **Figure 2**, participants were both more accurate (**Fig. 2a**) and faster (**Fig. 2b**) in trials with attention-directing versus neutral cues. This was corroborated statistically, both for accuracy (t(23) = 9.798, p = 1.123e-9, d = 2.0) and for reaction time (t(23) = -6.830, p = 5.772e-7, d = - 1.394).

**Figure 2.**
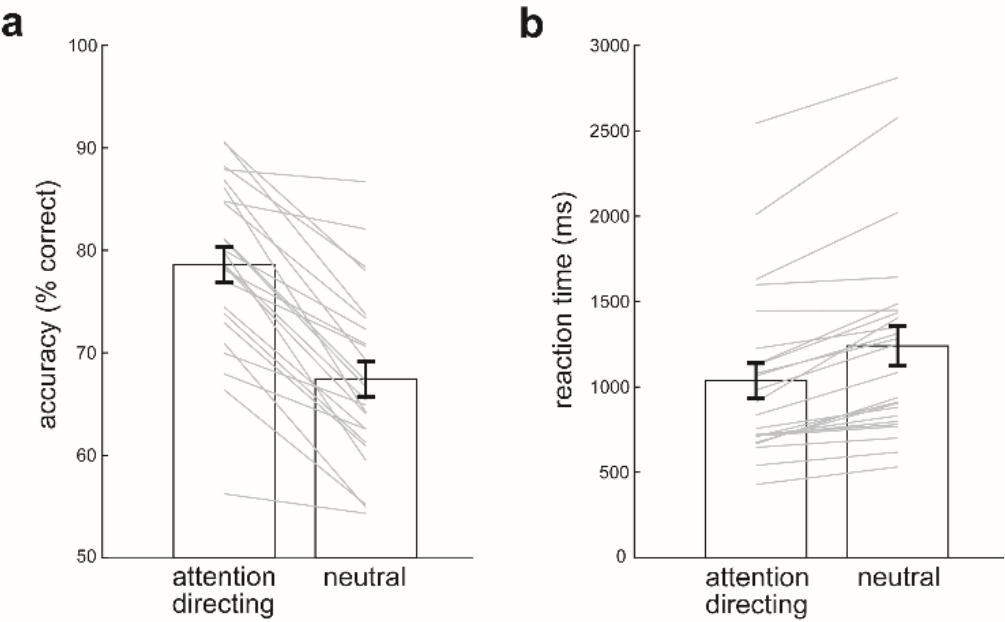
Behavioural performance confirms participants used the cue when possible. **a)** Task accuracy in trials with attention-directing and neutral cues. **b)** Reaction times in trials with attention-directing and neutral cues. Error bars indicate ± 1 SEM calculated across participants (n=24). Grey lines denote individual participants.

### Attentional cues during working memory lead to a robust spatial modulation in microsaccades

Having confirmed that participants used the informative (attention-directing) cues to improve performance, we next assessed how informative cues – that directed attention to memory items that had been presented to the left or right at encoding – modulated the direction of microsaccades. Building on our prior studies (de Vries et al., 2023; Liu et al., 2022; van Ede et al., 2019, 2020) as well as related studies deploying external covert-attention tasks (Corneil & Munoz, 2014; Engbert & Kliegl, 2003; Fernández et al., 2023; Hafed et al., 2011; Hafed & Clark, 2002; Lowet et al., 2018), we observed robust biasing of saccade directions as a function of whether cues directed attention to memory items that were presented to the left or to the right of fixation at encoding (**Fig 3a**). When statistically evaluating the full-time course, we observed two consecutive significant clusters (horizontal lines in **Fig. 3a**; cluster P values: 0.01 and 3.9996e-04). Likewise, when zooming in on the a-priori defined time window from 200-600 ms after the cue (based on (Liu et al., 2022)), we observed a highly robust modulation (**Fig 3b**; t(23)= 4.171, p = 0.0004, d = 0.851) with more saccades toward than away from the memorised location of the cued item.

**Figure 3.**
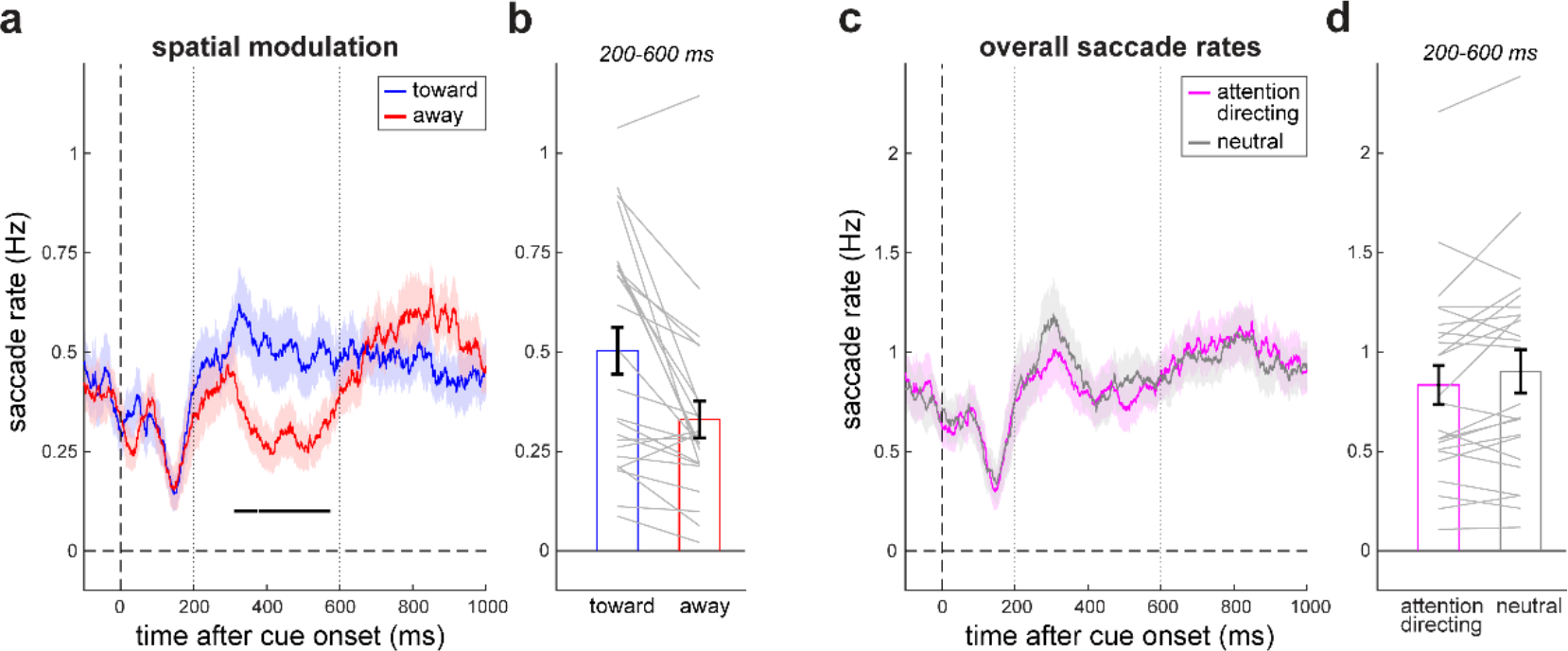
Internal selective attention modulates the direction of microsaccades without changing overall microsaccade rate. **a)** Saccade rates in trials with attention-directing cues as a function of time after cue onset for saccades in the direction of the memorised location of the cued memory item (toward) and in the opposite direction (away). Shadings indicate ± 1 SEM calculated across participants (n=24). The black horizontal lines denote significant clusters following cluster-based permutation analysis (Maris & Oostenveld, 2007). Dashed vertical lines indicate the a-priori defined time window of interest, from 200 to 600 ms after cue onset. **b)** Bar graph of toward and away saccade rates from panel a, averaged over the a-priori defined window from 200-600 ms after cue onset (based on (Liu et al., 2022)). Error bars indicate ± 1 SEM calculated across participants (n=24). Grey lines denote individual participants. **c)** Overall saccade rates as a function of time after cue onset following attention-directing and neutral cues. Note how saccade rates are than in panel a, given that overall saccade rates include both toward and away saccades. **d)** Bar graph of overall saccade rates in panel c, averaged over the a-priori defined window from 200-600 ms post cue onset.

To establish the nature of this spatial saccade modulation, we repeated the above analysis as a function of saccade size (**Fig. 4a**). As can be seen, the vast majority of saccades occurred in the microsaccade range, below 1 degree visual angle. This is perhaps not surprising given that in this time period of interest, there was nothing on the screen apart from the fixation dot. Critically, when directly comparing toward and away saccades (**Fig. 4a**, bottom panel), we also found that the spatial modulation was confined to the microsaccade range, replicating our previous findings (de Vries et al., 2023; Liu et al., 2022; van Ede et al., 2019).

**Figure 4.**
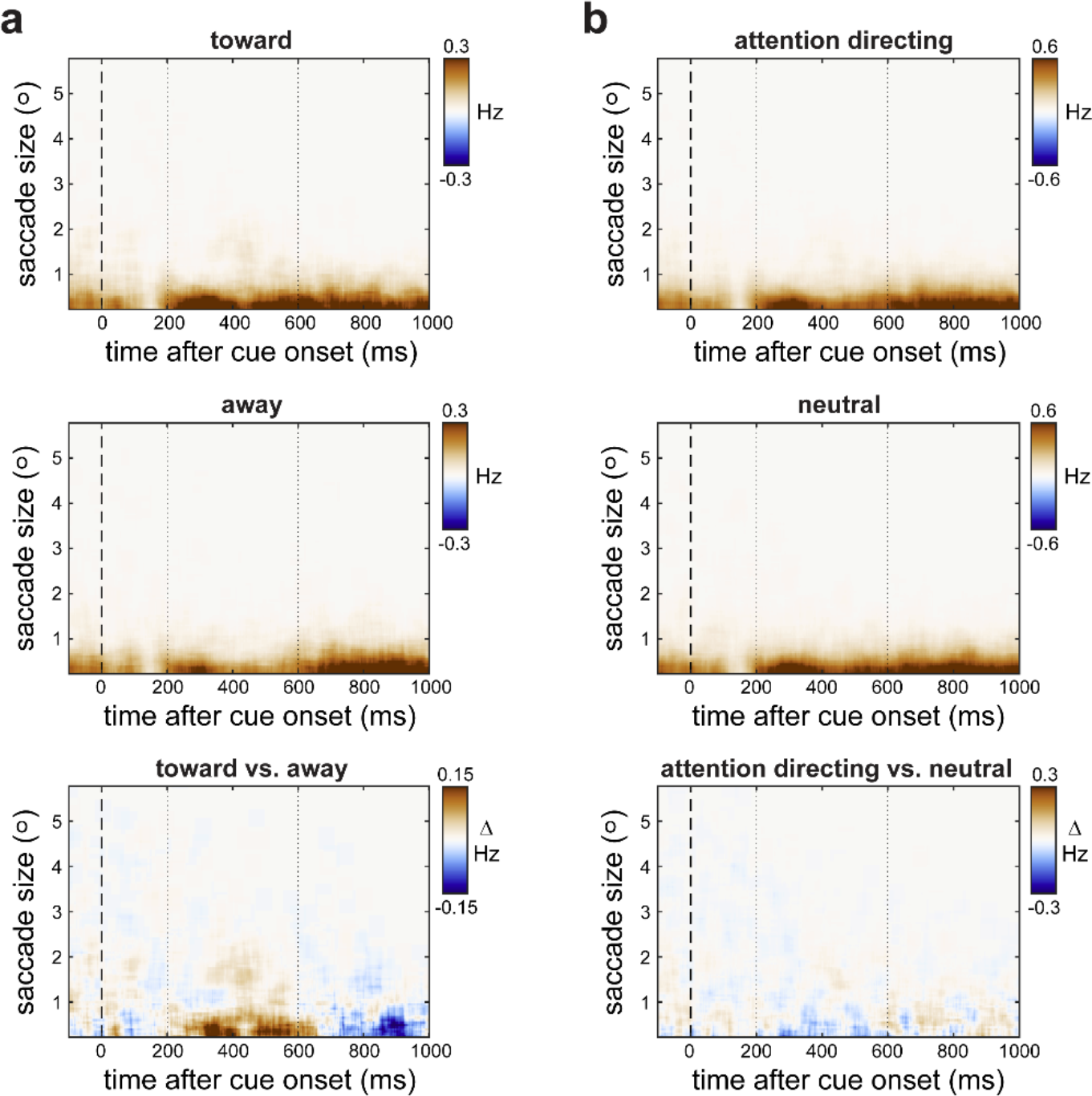
Attentional biasing of saccades is driven by saccades in the microsaccade range. **a)** Saccade rates as a function of saccade size (y axes) and time after attention-directing cues (x axes) for toward saccades (top), away saccades (middle), and their difference (toward minus away; bottom). This shows how the vast majority of saccades occur below 1 degree visual angle (i.e. in the microsaccade range) and how the directional bias (bottom) is also largely confined to the microsaccade range (as also in (Liu et al., 2022)). During encoding, items were centred at ± 5 degrees to the left and right of fixation. **b)** Overall saccade rates as a function of saccade size and time after cue onset for trials with attention-directing cues (top), neutral cues (middle), and their difference (attention-direction minus neutral; bottom).

In our task, the spatial biasing of microsaccades must reflect a shift of attention to the colour-matching memory item that was held within the spatial lay-out of working memory. It cannot reflect sensory processing of the cue or anticipation of the probe, as both cue and probe were always presented centrally (i.e., left and right were exclusively defined in the memorised visual space).

### The attentional modulation of microsaccades is not driven by the addition of new microsaccades

We finally turn to the central question of the current study: whether the above-described modulation of microsaccades is driven by the addition of new, attention-driven, microsaccades or whether this spatial modulation is driven by a directional biasing of ongoing microsaccades that would have been made anyway. For this, we compared overall microsaccade rates following informative, attention-directing, cues versus following neutral cues. Our logic was straightforward: if attention introduces new microsaccades then the overall rate should increase following attention-directing compared to neutral cues.

Overall saccade rates following attention-directing and neutral cues are shown in **Figure 3c-d**. In contrast to the above prediction, we found no evidence for an increase in overall microsaccade rate following attention-directing cues compared to following neutral cues (that were matched in terms of bottom-up sensory stimulation). In fact, if anything, we found a slight decrease in overall microsaccade rate following attention-directing cues, though this did not reach significance – neither when considering the full-time axis (no significant clusters; **Fig. 3c**), nor when zooming in on the a-priori defined window of interest (**Fig. 3d**; t(23) = -1.994, p = 0.058, d = -0.407). This implies attention shifts do not generate new microsaccades.

To complement this main result, **Figure 4b** shows the relevant data as a function of saccade size. This confirmed a similar prevalence of saccades below 1 degree following both attention-directing and neutral cues, with no evidence for more saccades in this microsaccade range following attention-directing cues (**Fig. 4b**, lower panel), despite the clear spatial modulation that we observed following these cues.

## DISCUSSION

We observed robust modulations in microsaccade direction, with more microsaccades toward versus away from the memorised location of a cued visual memorandum (replicating our previous work, e.g., (Liu et al., 2022; van Ede et al., 2019)). Our aim was to assess whether this spatial modulation is driven by the addition of new microsaccades that are triggered directly by a spatial shift in attention. To this end, we compared overall microsaccades rates following attention-directing cues to a control condition with neutral cues that did not invite any shift of attention, but that were matched in sensory properties otherwise. Our data showed no evidence for an increase in overall microsaccade rate following attention-directing cues (if anything we observed a slight, albeit non-significant decrease). This lack of a rate increase in the face of a clear directional modulation implies that the attentional modulation of microsaccades is not driven by the injection of “new” microsaccades. Instead, these data suggest that attention merely biases the direction of ongoing microsaccades that would have been made whether or not attention was also shifted. In other words, shifting attention does not change the probability that a microsaccade will occur, but it does change the probability where a microsaccade will go – *if* one will be made.

By studying microsaccades as an accessible peripheral signature of upstream oculomotor brain circuitry, our findings have implications for our understanding of the links between attention and the oculomotor system. For example, it has previously been established that the superior colliculus, besides regulating saccades and microsaccades, may also play a key role in shifting covert attention (e.g., (Krauzlis et al., 2013; Lovejoy & Krauzlis, 2010; Muller et al., 2005)). Our data are consistent with this, and tentatively suggest that attention shifts may not necessarily add to ongoing activity within the superior colliculus, as evidenced by the absence of an increase in overall microsaccade. Instead, we speculate based on our data that attention may re-balance activity of the pool of superior-colliculus neurons (whose overall activity levels and excitatory/inhibitory balance may be normalised, for example, via a divisive normalisation mechanism; (Carandini & Heeger, 2012; Reynolds & Heeger, 2009)). Such re-balancing would predict that the distribution of activity may vary depending on attention, but not the total amount of activity – consistent with our finding of not more microsaccades, but instead the same number of microsaccades that go more in the attended direction.

Previous work has suggested that microsaccades are correlated with, but not necessary for attentional shifts (e.g., (Horowitz et al., 2007; Liu et al., 2022; Yu et al., 2022)). Our findings are consistent with, and help to appreciate, this probabilistic nature of this link between microsaccades and selective attention. Because attention shifts themselves do not trigger microsaccades, it is possible to have attention shifts without a peripheral trace in the form of a microsaccade. Instead, only when attention shifts are made in the presence of an (already planned) microsaccade, will we observe a correlation between microsaccade direction and the direction of the covert or internal shift of attention. It is noteworthy, however, that we here studied microsaccades when participants are explicitly cued to voluntarily shift attention. Complementary work has shown how spontaneous microsaccades – made in the absence of volitional shifts of attention – may themselves trigger performance and neural modulations that are typically associated with attention shifts (Hafed, 2013; Shelchkova & Poletti, 2020; Yuval-Greenberg et al., 2014). Whether and how attention can become decoupled from such spontaneous microsaccades, or whether attention may inevitably follow in the case of spontaneous microsaccades, remains an interesting question not addressed by the current study.

A recent study (Willett & Mayo, 2023) found little to no evidence for a directional biasing of microsaccades to an attended visual stimulus, despite clear behavioural and neural benefits of attention. A critical difference with our study is that the authors did not consider *shifts* of attention following a cue, but rather *sustaining* attention to either of two targets that remained fixed throughout a block of trials. It is conceivable that microsaccade biases may be particularly sensitive to shifts of covert selective attention, without necessarily also tracking the process of sustaining covert visual attention after this initial shift (van Ede, 2023). Another set of complementary studies focused on microsaccades following exogenous capture of attention to a peripheral cue. These studies have typically reported microsaccades biases that look away from the cued location (e.g., (Engbert, 2012; Laubrock et al., 2005)), rather than the observation of more toward microsaccades that we reported here, and that previous studies also reported following voluntary attention cues. Whether such microsaccades biases that occur in the opposite direction also reflect biasing of ongoing microsaccades or rather the addition of new microsaccades remains an interesting question for future work.

At least two early studies on microsaccade biases by attention also included neutral cues (Engbert & Kliegl, 2003; Laubrock et al., 2005). While their data thus allowed the same key comparison as we targeted here, there are several relevant differences. The most critical difference is that in these studies the employed endogenous cues showed only weak directional biasing of microsaccades (Laubrock et al., 2005). In comparison, here, we observed a clear bias; building on our prior work using the same overall task set-up (e.g., (Liu et al., 2022; van Ede et al., 2019, 2020, 2021)). It was only in the presence of this clear spatial modulation that our question was of interest: whether we could “explain” this spatial modulation by the addition of new microsaccades. In addition, in aforementioned work, the authors did not fully match informative (attention-directing) and neutral cues. In contrast, we always used the same colour cues, and counterbalanced whether specific colours were attention-directing or neutral. Finally, we here uniquely compared microsaccades following attention-directing versus neutral cues in the context of an internal selective attention task in which attention was directed internally to the contents of working memory – a task that yielded robust spatial modulations that were a pre-requisite for addressing our central question.

In conclusion, we report a clear biasing of microsaccades by the direction of internal attention shifts and reveal how this microsaccade modulation must be attributed to a spatial biasing of ongoing microsaccades rather than by the addition of new microsaccades that are triggered by the attention shift itself. This helps to explain the probabilistic nature of the link between microsaccades and attention and provides relevant context for future work delineating the links between attention, microsaccades, and upstream oculomotor brain circuitry.

## ACKNOWLEDGEMENTS

This research was supported by an ERC Starting Grant from the European Research Council (MEMTICIPATION, 850636) and an NWO Vidi grant by the Dutch Research Council (grant number 14721) to F.v.E. The authors also wish to thank Eelke de Vries for his valuable comments on the manuscript.

## AUTHOR CONTRIBUTIONS

B.L and F.v.E designed the experiment; B.L. programmed the task; Z-S.A acquired the data; B.L and F.v.E analysed and interpreted the data; B.L and F.v.E drafted and revised the manuscript.

